# Bitter taste receptor-stimulated nitric oxide innate immune responses are reduced by loss of CFTR function in nasal epithelial cells and macrophages

**DOI:** 10.1101/2022.09.21.508717

**Authors:** Ryan M. Carey, James N. Palmer, Nithin D. Adappa, Robert J. Lee

**Affiliations:** Department of Otorhinolaryngology—Head and Neck Surgery; Department of Physiology, University of Pennsylvania Perelman School of Medicine

**Keywords:** Chronic rhinosinusitis, cystic fibrosis, mucociliary clearance, phagocytosis, nitric oxide

## Abstract

Bitter taste receptors (T2Rs) are G protein-coupled receptors identified on the tongue but expressed all over the body, including in airway cilia and macrophages, where T2Rs serve an immune role. T2R isoforms detect bitter metabolites (quinolones and acyl-homoserine lactones) secreted by gram negative bacteria, including *Pseudomonas aeruginosa*, a major pathogen in cystic fibrosis (CF). T2R activation by bitter bacterial products triggers calcium-dependent nitric oxide (NO) production. In airway cells, the NO increases mucociliary clearance and has direct antibacterial properties. In macrophages, the same pathway enhances phagocytosis. Because prior studies linked CF with reduced NO, we hypothesized that CF cells may have reduced T2R/NO responses. Using qPCR, immunofluorescence, and live cell imaging of air-liquid interface cultures, we found primary nasal epithelial cells from both CF and non-CF patients exhibited similar T2R expression, localization, and calcium signals. However, CF cells exhibited reduced NO production also observed in immortalized CFBE41o-CF cells and non-CF 16HBE cells CRISPR modified with CF-causing mutations in the CF transmembrane conductance regulator (CFTR). NO was restored by VX-770/VX-809 corrector/potentiator pre-treatment, suggesting reduced NO in CF cells is due to loss of CFTR function. In nasal cells, reduced NO correlated with reduced ciliary and antibacterial responses. In primary human macrophages, inhibition of CFTR reduced NO production and phagocytosis during T2R stimulation. Together, these data suggest an intrinsic deficiency in T2R/NO signaling caused by loss of CFTR function that may contribute to intrinsic susceptibilities of CF patients to *P. aeruginosa* and other gram-negative bacteria that activate T2Rs.

## Introduction

Cystic fibrosis (CF) is a common lethal recessive genetic disease characterized in part by reduced airway surface liquid volume, overly thick airway surface mucus, and impaired mucociliary clearance (1, 2). This is caused by defective fluid secretion and/or absorption upon loss of function of the CF transmembrane conductance regulator (CFTR). CFTR is an anion channel (3) that supports Cl^−^, HCO_3_^−^, and fluid secretion in airway submucosal gland and surface epithelial cells (4-8). Because mucociliary clearance is the airway’s most important physical defense and depends on proper fluid volume and mucus rheology (9, 10), lack of proper CFTR function and failure to properly clear inhaled/inspired bacteria results in increased incidence of both upper (9, 11-13) and lower (14, 15) respiratory infections. CF was, until recently, almost always fatal without a lung transplant. Small molecule ion channel modulators that restore CFTR function are emerging as potentially highly effective therapies for CF (3). Nonetheless, some patients (e.g., those with CFTR premature stop mutations like G542X) cannot yet benefit from small molecule therapies. Additionally, a myriad of cellular defects have been suggested to occur due to lack proper CFTR function, including impaired macrophage function (16-18) and/or kinase signaling (19), either secondary to ion transport or independent of ion transport, possibly due to CFTR scaffolding regulation of cellular signaling. Understanding other cellular processes affected by CFTR mutations as well as if/how modulators restore these processes is important to optimizing therapies for CF lung disease and CF-related chronic rhinosinusitis (CRS).

Leveraging endogenous immune receptors to stimulate or enhance innate defensive responses instead of using single agent antibiotics could reduce the selective pressure responsible for antibiotic-resistant pathogens, an important issue for CF patients (14, 20). Taste family 2 receptors (T2Rs) are G protein-coupled receptors (GPCRs) responsible for bitter taste on the tongue, which are expressed within bronchial and nasal cilia. These T2Rs detect bacterial products, including *P. aeruginosa* acyl-homoserine lactone (AHL) (21) and quinolone (22) metabolites to activate local and rapid (within minutes) defensive responses involving nitric oxide production. NO generated in the nose is important for airway immunity because it increases mucociliary clearance through activation of guanylyl cyclase-dependent cGMP production and protein kinase G (PKG) phosphorylation of cilia proteins (23). NO also directly kills or inactivates pathogens (24). NO damages cell walls and DNA of bacteria (25-29). Replication of many respiratory viruses is also NO-sensitive, including influenza, parainfluenza, rhinovirus (30), and SARS-COV1 & 2 (31-34).

While there are 25 T2R isoforms on the tongue, several (T2Rs 4, 14, 16, and 38) are expressed in nasal cilia. These T2Rs stimulate Ca^2+^-dependent NOS—specifically the same eNOS isoform implicated in reduced NO production in CF endothelial cells (35); eNOS is also localized to cilia (36-38). T2R activation of eNOS in cilia produces NO to increase cilia beating through protein kinase G and more directly kill bacteria (21, 39, 40). T2Rs are also expressed in immune cells like macrophages (41-43) and neutrophils (44, 45). We found that T2Rs in macrophages enhance phagocytosis in response to bacterial metabolites, also through Ca^2+^ and NO signaling (42), underscoring the importance of T2R-stimulated NO generation.

Clinical studies in non-CF patients support the *in vivo* importance of NO-producing T2Rs in immune defense in chronic rhinosinusitis (CRS). *TAS2R38* is the gene encoding a highly expressed T2R isoform in cilia, T2R38. *TAS2R38* has two polymorphisms with a Mendelian distribution in most populations (39, 46-48). The polymorphisms result in either a proline-alanine-valine (PAV) or an alanine-valine-isoleucine (AVI) at three positions in T2R38. The PAV T2R38 variant is functional, while the AVI variant is not, based on taste and *in vitro* responses to T2R38-specific agonist phenylthiocarbamide (PTC) (49). AVI/AVI nasal cells (non-functional T2R38) produce only ∼10% of the NO produced by PAV/PAV cells in response to PTC, bacterial lactones, or *P. aeruginosa* conditioned media (39). Patients homozygous for AVI *TAS2R38* (non-functional T2R38; ∼25% of the population) are more susceptible to gram-negative upper respiratory infection (39) and CRS (46, 48). AVI/AVI individuals have worse sinus surgical outcomes than the ∼25% of the population homozygous for PAV (functional) T2R38 (50). This was supported by Canadian (51) and Polish (52) genome-wide association studies of *TAS2R38* with CRS, an Australian study showing AVI/AVI T2R38 homozygotes have increased sinonasal bacterial load (53), and others. These data support T2Rs as an early warning arm of innate immunity important *in vivo* in the upper respiratory tract.

*TAS2R38* genotype correlates with sinonasal quality of life in CF CRS patients (47), based on SNOT-22 scoring, a standard metric in CRS research (54). CF individuals with PAV/PAV (functional T2R38) genotype had lower SNOT-22 scores (n = 49, *p* <0.05), indicating lower symptom burden. Specifically, nasal symptoms were less severe in PAV/PAV CF patients compared to individuals with other genotypes (n = 47, *p* <0.05). However, other studies report *TAS2R38* genotype does not correlate with need for sinus surgery (55) nor with *P. aeruginosa* lung infection (56). A more recent study concluded that *TAS2R38* is a modifier of CF (57), with PAV (functional T2R38) allele frequency lower in CF patients colonized with *P. aeruginosa* before age 14.

We hypothesized that the somewhat subtle effects of TAS2R38 genotype on CF disease progression might reflect a general defect in this pathway in CF cells. While there are conflicting reports about NO in CF airways, some studies have suggested reduced airway NO in CF patients correlated with *Pseudomonas aeruginosa* infection (58-61). Defects in CFTR function have been proposed to reduce kinase-dependent activation of eNOS in CF endothelial cells downstream of Akt kinase activation via sheer stress. If T2R-activated NO signaling (occurring via the same eNOS enzyme in airway cells, macrophages, and endothelial cells) is reduced in CF, this may allow *P. aeruginosa*, which produces AHLs and quinolones that activate T2Rs, to thrive in CF airways. *P. aeruginosa* is more sensitive to NO killing than many other airway pathogens, e.g., *Staphylococcus aureus* (25).

To test this, we grew primary nasal cells isolated from CF and non-CF patients and genotyped them for *TAS2R38*. Cells were differentiated at air-liquid interface, a gold-standard airway epithelial model (62), and T2R localization and function was examined using immunofluorescence microscopy, live cell imaging, and microbiological assays. We also tested T2R function in immortalized cell line-based isogenic CFTR mutation models. The data below suggest reduction in T2R-mediated NO generation with lack of proper CFTR function, suggesting a novel intrinsic reduction of innate immunity in CF cells that is ion transport-independent.

## Materials and Methods

### Cell culture

Immortalized cells were grown in minimal essential media (MEM; ThermoFisher Scientific) plus 1x cell culture penicillin/streptomycin (Gibco) and 10% FetalPlex serum substitute (Gemini Biosciences). Parental CFBE41o-cells (F508del CFTR homozygous (63) and CFBE41o-cells stably expressing either Wt CFTR or F508del CFTR with puromycin resistance (64, 65) were a gift from Dr. R.C. Rubenstein (Division of Allergy, Immunology and Pulmonary Medicine, Washington University in St. Louis). CRISPR-modified 16HBE14o-cells homozygous for G542X or F508del CFTR (66) were obtained from the Cystic Fibrosis Foundation under a materials transfer agreement. Identity was confirmed by in-house sequencing (University of Pennsylvania Department of Genetics Next-Generation Sequencing Core).

For air-liquid interface (ALI) cultures, 16HBE14o- or CFBE41o-cells were seeded onto collagen-coated 0.33 cm^2^ transwell filters and grown to confluence for 5 days before apical air exposure, as described previously (67, 68). Upon air exposure, basolateral media was changed to a differentiation media containing 1:1 Lonza bronchial epithelial cell basal media (BEBM):DMEM plus Lonza singlequot supplements (0.5 ng/ml hEGF, 5 ng/ml epinephrine, 0.13 mg/ml BPE, 0.5 ng/ml hydrocortisone, 5 ng/ml insulin, 6.5 ng/ml triiodothyronine, and 0.5 ng/ml transferrin, 0.1 nM retinoic acid) supplemented with 1x penicillin/streptomycin and 2% NuSerum (BD Biosciences, San Jose, CA) as described (39). Cells were fed from the basolateral side only with differentiation media for ∼21 days before use.

Primary human nasal epithelial cells were isolated from surgical specimens according to The University of Pennsylvania guidelines regarding use of residual clinical material. Tissue was from patients undergoing sinonasal surgery at the Hospital of the University of Pennsylvania under institutional review board approval (#800614) with written informed consent in accordance with the U.S. Department of Health and Human Services code of federal regulation Title 45 CFR 46.116 and the Declaration of Helsinki. Inclusion criteria were patients ≥18 years of age requiring surgery for sinonasal disease or trans-nasal approaches to the skull base. Exclusion criteria included systemic inheritable disease (eg, granulomatosis with polyangiitis, systemic immunodeficiences) or use of antibiotics, oral corticosteroids, or anti-biologics (e.g. Xolair) within one month of surgery. Vulnerable populations (patients ≤18 years of age, pregnant women, and cognitively impaired persons) were not included.

Tissue was transported to the lab on ice in saline. Mucosal tissue was immediately removed for enzymatic dissociation. Cells were grown to confluence in proliferation medium for 7 days, dissociated, and seeded at high density on Corning transwells coated with type I bovine collage, fibronectin, and bovine serum albumin (22, 39, 68, 69). Culture medium was removed from the upper compartment the next day, and basolateral media was changed to differentiation medium as described above. Cultures were genotyped for *TAS2R38* PAV or AVI polymorphims (49, 70) as described (22, 39, 68). PAV/AVI heterozygote cells were used in experiments with T2R14 agonists like quinine, apigenin, or diphenhydramine, as we have previously found these responese to be independent of *TAS2R38* genotype (68, 71, 72).

CF cells used here were identified as homozygous for phenyalanine 508 deletion (F508del/F508del) or to have one copy of F508del plus one copy of a minimally functional CFTR variant (e.g., G542X) based on their patient medical record. Because CF CRS patients typically undergo surgery at a younger age than the average CRS patient, and because age might be a factor in airway cell NO production, we used cells from a subset of younger CRS patients (mean age at surgery 35 ± 2 years) to match the age of our CF CRS patients (mean age at surgery 33 ± 3 years, *p* = 0.56 by Student’s *t* test). The two patient populations are shown in **Supplemental Table 1**.

Primary human monocyte-derived macrophages were cultured as previously reported (73, 74). De-identified human monocytes isolated from healthy apheresis donors were obtained by the University of Pennsylvania Human Immunology core. M0 macrophages were differentiated by adherence culture in RPMI 1640 + 10% human serum + 1x pen/strep for 12 days in 8-well chamber slides (CellVis) in high glucose RPMI2650 supplemented with 1x cell culture Pen/Strep and 10% human serum. Our prior studies suggest no differences in T2R responses among macrophages differentiated by adherence alone or by adherence plus M-CSF (42), and thus adherence alone was used for these studies.

### Ca^2+^ and NO imaging

Fura-2 (Ca^2+^ indicator dye) and DAF-FM (NO indicator dye) were imaged as previously described (68, 73). Briefly, fura-2 was imaged using MetaFluor (Molecular Devices, Sunnyvale, CA USA) and dual excitation filter set on an IX-83 microscope (10x 0.4 NA PlanApo objective) equipped with a fluorescence xenon lamp (Sutter Lambda LS, Sutter Instruments, Novato, CA USA), excitation and emission filter wheels (Sutter Lambda 10-2), and Orca Flash 4.0 sCMOS camera (Hamamatsu, Tokyo, Japan). DAF-FM was imaged on a TS100 microscope (10x 0.3 NA PlanFluor objective; Nikon, Tokyo, Japan) with GFP filter set, XCite 110 LED (Excelitas Technologies, Waltham MA USA), and Retiga R1 Camera (Teledyne QImaging, Surrey, BC, Canada). DAF-FM time course images were acquired using Micromanager variant of ImageJ (75). Primary human ALIs were loaded for 90 min with 10 µM DAF-FM-diacetate or fura-2 acetoxymethyl ester (AM) in 20 mM HEPES-buffered HBSS supplemented with 1x MEM amino acids on the basolateral side and glucose free HEPES-buffered HBSS (no amino acids) on the apical side, followed by washing with the same buffer (22, 39, 69). Compounds were added to the apical side in glucose-free HBSS. Macrophages were loaded with 5 µM fura-2-AM or DAF-FM DA for 45 min as previously described (73, 74) and imaged with 20x 0.75 NA PlanApo objective.

Cells were stimulated with 10 µg/ml SC79 that was made from 10 mg/ml SC79 stock in DMSO. PTC was made as 1 M stock in DMSO, 3oxoC12HSL as 100 mM stock in DMSO, apigenin as 100 mM stock in DMSO. Control solutions were HBSS + 0.1% or 0.2% DMSO vehicle control as appropriate. No effects of DMSO alone were observed, as shown in figures below. Quinine, denatonium benzoate, and sodium benzoate were dissolved directly in HBSS.

### Measurement of ciliary beat frequency (CBF)

Whole-field CBF was imaged at 120 frames per second using a Basler A602 camera and Nikon TS-100 microscope (40x long working distance objective) at ∼26-28 °C in a custom glass bottom chamber. Experiments utilized Dulbecco’s PBS (+ 1.8 mM Ca^2+^) on the apical side and 20 mM HEPES-buffered Hank’s Balanced Salt Solution supplemented with 1× MEM vitamins and amino acids on the basolateral side. Data were analyzed using the Sisson-Ammons Video Analysis system and normalized to baseline CBF as previously described (4, 39, 76-79).

### Immunofluorescence (IF) microscopy

IF was carried out as previously described (22, 39, 68), with ALI cultures fixed at room temperature in 4% formaldehyde for 20 min, followed by blocking and permeabilization for 1 hour at 4°C in Dulbecco’s phosphate buffered saline (DPBS) containing 5% normal donkey serum (NDS), 1% bovine serum albumin (BSA), 0.2% saponin, and 0.3% Triton X-100. After three washes in DPBS, primary antibody incubation (1:100 for anti-T2R antibodies, 1:250 for tubulin antibody) was performed overnight at 4°C in DPBS containing 5% NDS, 1% BSA, and 0.2 % saponin. Subsequent incubation with AlexaFluor (AF)-conjugated donkey anti-mouse and anti-rabbit secondary antibodies (1:1000) was done for 2 hours at 4°C. Transwell filters were then removed from the plastic mounting ring and mounted with Fluoroshield with DAPI (Abcam; Cambridge, MA USA)). For co-staining of T2R14 and T2R38, primary antibodies were labeled directly using Zenon antibody labeling kits (Thermo Fisher Scientific) for AF546 or AF647 as described (22, 39, 68). Images of ALIs were taken on an Olympus Fluoview confocal system with IX-73 microscope and 60x (1.4 NA) objective and analyzed in FIJI (80). Images of submerged H441 cells were taken on an Olympus IX-83 microscope with 60x (1.4 NA) objective using Metamorph. Anti-T2R38 (ab130503; rabbit polyclonal) and anti-beta-tubulin IV (ab11315; mouse monoclonal) antibodies were from Abcam. Anti-T2R14 (PA5-39710; rabbit polyclonal) primary antibody, Anti-T2R46 (rabbit polyclonal; OSR00137W), and conjugated secondary antibodies (donkey anti-rabbit AlexaFluor 546 and donkey anti-mouse AlexaFluor 488) were from ThermoFisher Scientific. T2R38 antibody (Cat #AP59054) and C-terminal peptide (Cat #BP16929b) were from Abcepta. Blocking peptide was incubated with primary antibody at 10:1 molar ratio for 4 hrs at 4°C prior to use. T2R14 antibody (LS-C413403-100) and blocking peptide (LS-E44266-1) were from LSBio. Immunofluorescence images were analyzed in FIJI (80) using only linear adjustments (min and max), set equally between images obtained at identical microscope settings (exposure, objective, binning, etc.).

### Bacteria culture

*P. aeruginosa* strains PAO-1 (ATCC 15692) and clinical CRS-isolate strain P11006 (obtained from Drs. N. Cohen and L. Chandler, Philadelphia VA Medical Center) (72) were grown in LB media (Gibco). For anti-bacterial assays, P. aeruginosa were grown to OD 0.1 and resuspended in 50% saline containing 1 mM HEPES and 0.5 mM glucose, pH 6.5. Nasal ALIs were washed 24 hrs prior with antibiotic-free Ham’s F12K media (ThermoFisher Scientific) on the basolateral side. 30 uL of bacteria solution was placed on the apical side of the ALI for 10 min, followed by aspiration of bulk ASL fluid. After 2 hrs at 37 °C, remaining bacteria were removed from the ALI culture by washing followed by life-dead staining with SYTO9 (live) and propidium iodide (dead) with BacLight Bacterial Viability Kit (ThermoFisher Scientific). Control experiments were performed similarly using transwell filters with no cells and bacteria in solution ± 10 µg/ml colistin sulfate. Green (live)/red (dead) ratio was quantified in a Spark 10M (Tecan, Mannedorf, Switzerland) at 485 nm excitation and 530 nm and 620 nm emission.

### Phagocytosis assays

Phagocytosis assay (as descried (73, 74)) were carried out by incubating macrophages with heat-killed FITC-labeled *Escherichia coli* strain K-12 bioparticles (ThermoFisher Scientific Vybrant phagocytosis assay kit) at 250 µg/ml in phenol red-free, low glucose DMEM (15 min, 37°C) followed by immediate recording of fluorescence from living cells after quenching extracellular FITC with trypan blue per the manufacturer’s instructions. Fluorescence was recorded on a Spark 10M plate reader (Tecan) with 485 nm excitation and 535 nm emission.

### Data analysis and statistics

Data were analyzed in Excel (Microsoft) and/or Prism (GraphPad software, La Jolla, CA). All data in bar graphs are shown as mean ± SEM. Multiple comparisons were made in Prism using one-way ANOVA with Bonferroni (pre-selected pairwise comparisons), Tukey-Kramer (comparing all values), or Dunnett’s (comparing to control value) post-tests; *p* <0.05 was considered statistically significant. Asterisks (* and **) indicate *p* <0.05 and *p* <0.01, respectively. Data used to construct graphs are available upon request.

## Results

We examined primary nasal epithelial cells grown and differentiated at air-liquid interface (ALI), which form motile cilia (**Fig 1A**) that also express bitter receptors like T238 (**Fig 1B-C**) (39), T2R4, T2R14, and T2R16 (**Fig 1D**). Over the course of ALI differentiation, there was no significant difference between TAS2R expression in F508del/F508del CF vs non-CF cells by qPCR (**Fig 1D**). Co-staining for T2R38 and T2R14 suggested that both T2R isoforms were cilia-localized in both CF and non-CF cells (**Supplemental Fig 1A-C**), and co-localization was similar when measured by Pearson’s correlation coefficient (**Supplemental Fig 1D**) or antibody-based FRET efficiency calculations (**Supplemental Fig 2A-C**). Thus, both T2R expression and localization appear to be similar in CF and non-CF cells.

**Figure 1.**
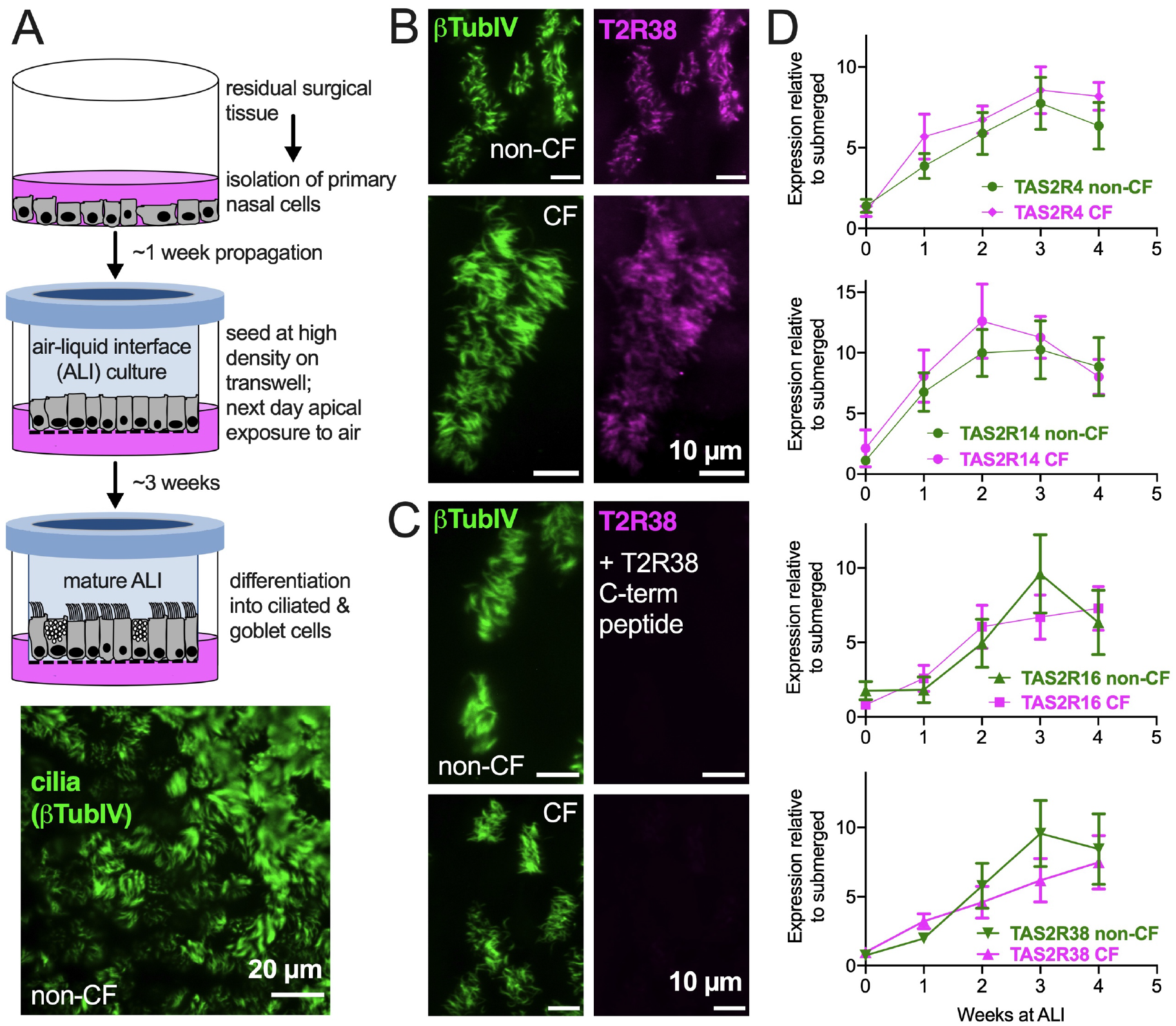
Bitter taste receptor (T2R) transcript expression in differentiated CF and non-CF primary nasal cells. **(A)** Air-liquid interface (ALI) culture model for differentiation of primary nasal epithelial cells (top) results in formation of motile cilia (bottom, stained with antibody against β-tubulin IV). **(B-C)** Nasal ALI cultures in differentiated CF (top panels) and non-CF primary nasal cells (bottom panels) express cilia-localized T238, evidenced by co-localization of β-tubulin IV. *B* shows immunofluorescence of C-terminal directed primary antibody while *C* shows immunofluorescence after primary antibody was incubated with 10-fold molar excess T2R38 C-terminal peptide prior to application on cells. Results representative of 3 independent experiments using cells from 3 patients. **(D)** Taqman qPCR for *TAS2R4, TAS2R14, TAS2R16*, and *TAS2R38* genes encoding cilia-localized T2R4, 14, 16 and 38 revealed no differences between CF and non-CF cells over the course of four weeks of mucociliary differentiation. Expression is normalized to housekeeping gene UBC. Results are mean ± SEM from 5 independent experiments using cells from 5 non-CF and 5 homozygous F508del CF patients. No significant differences determined by ANOVA.

**Figure 2.**
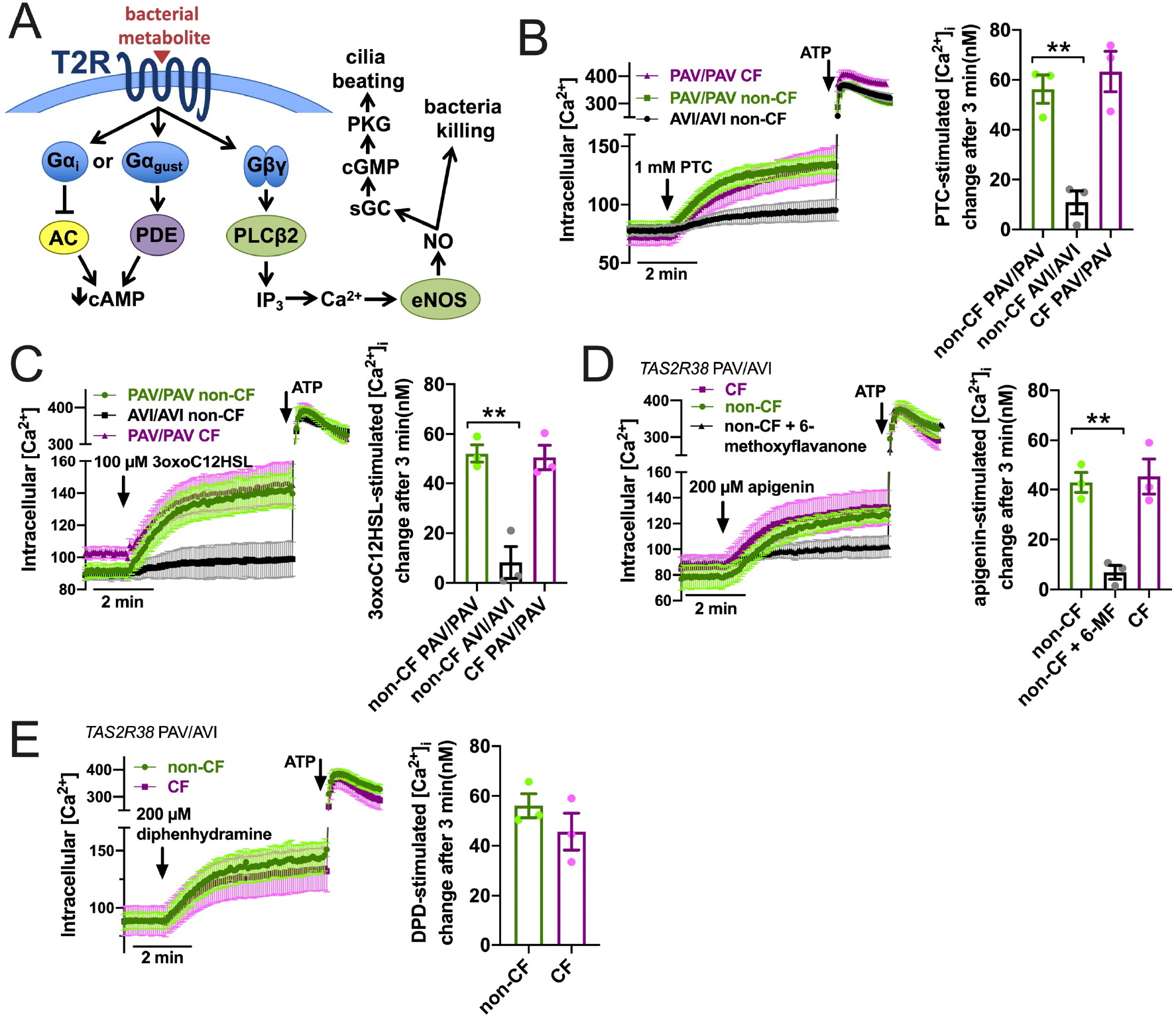
Ca^2+^ responses to T2R agonists were not different in CF vs. non-CF cells. **(A**) While the Gα arm of the T2R pathway typically lowers cAMP via Gα gustducin increase in phosphodiesterase (PDE) activity or Gαi lowering of adenylyl cyclase (AC) (24), T2R activation results in elevation of Ca^2+^ downstream of Gβγ-stimulated phospholipase C (PLC) to generate inositol trisphosphate (IP_3_), activating the IP_3_ receptor to release intracellular Ca^2+^ to activate endothelial nitric oxide synthase (eNOS) to produce nitric oxide (NO). The NO can both directly kill bacteria as well as activate soluble guanylyl cyclase (sGC) to increase cyclic GMP (cGMP) to activate protein kinase G (PKG), which phosphorylates ciliary proteins to increase ciliary beating. **(B)** Left, traces of Ca^2+^ indicator dye Fluo-4 in primary nasal air-liquid interface cultures (ALIs) genotyped as *TAS2R38* PAV/PAV (homozygous functional) CF or non-CF as well as non-CF *TAS2R38* AVI/AVI (homozygous non-functional) during stimulation with T2R38 agonist phenylthiocarbamide (PTC). Right, bar graph showing peak PTC-stimulated Ca2+ response. No significant difference between non-CF and CF PAV/PAV, but non-CF AVI/AVI was significantly lower than non-CF PAV/PAV by one-way ANOVA with Bonferroni posttest; \*\**p*<0.01. **(C)** Traces and bar graph similar to *B* but with cells stimulated with T2R38 agonist 3oxoC12HSL. No significant difference between non-CF and CF PAV/PAV, but non-CF AVI/AVI was significantly lower than non-CF PAV/PAV by one-way ANOVA with Bonferroni posttest; \*\**p*<0.01. **(D)** Traces (left) and bar graph (right) showing Fluo-4 Ca^2+^ responses in CF and non-CF ALIs genotyped as AVI/PAV *TAS2R38* during stimulation with T2R14 agonist apigenin. While there was no difference between CF and non-CF ALIs during apigenin stimulation, pre-incubation with T2R14 antagonist 6-methoxyflavanone (6-MF) during Fluo-4 dye loading reduced the apigenin response in non-CF cells; **p<0.01 by one way ANOVA with Bonferroni posttest. 0.1% DMSO was added to other conditions during loading as vehicle control for 6-MF. **(E)** Traces and bar graph similar to *D* but with T2R14 agonist diphenhydramine. No significant difference between CF and non-CF cells by Student’s t test. In all traces, 100 µM ATP (purinergic receptor agonist) is used as a positive control.

T2R activation results in Ca^2+^ elevation (**Fig 2A**). We measured Ca^2+^ responses downstream of T2R38 agonist PTC (49) using Ca^2+^ indicator fura-2. Responses to PTC were observed in non-CF cells genotyped for homozygous functional T2R38 (PAV variant) but not in non-CF cells genotyped for homozygous non-functional T2R38 (AVI variant) (**Fig 2B**). CF cells genotyped as PAV/PAV responded similarly to PAV/PAV non-CF cells (**Fig 2B**). We previously determined that Ca^2+^ responses to *P. aeruginosa* AHL 3-oxo-dodecanoyl-homoserine lactone (3oxoC12HSL (81)) in nasal ALIs is also dependent on T2R38. Like PTC, non-CF PAV/PAV (functional T2R38) cells responded to 3oxoC12HSL while non-CF AVI/AVI (non-functional T2R38) cells did not (**Fig 2C**). Also, like PTC, CF PAV/PAV and non-CF PAV/PAV cells responded similarly (**Fig 2C**). We also saw similar Ca^2+^ responses to T2R14 agonist apigenin (68) between CF and non-CF cells (**Fig 2D**). Responses in non-CF cells were blocked by antagonist 6-methoxyflavanone (82), supporting activation of T2R14 (**Fig 2D**). Finally, we noted similar Ca^2+^ responses to T2R14 agonist diphenhydramine (DPD) between CF and non-CF cells (**Fig 2E**).

Downstream of T2R Ca^2+^ signaling is NO production (**Fig 2A**). This occurs via endothelial nitric oxide synthase (eNOS) localized to the base of the cilia. Confirming this, we used a published siRNA protocol for primary ALIs (83) and found that eNOS siRNA but not nNOS siRNA reduced T2R Ca^2+^ responses (**Fig 3A**). We also noted that, despite nearly identical Ca^2+^ responses, F508del/F508del CF ALI cultures produced less NO in response to T2R agonists PTC, 3oxoC12HSL, quinine, apigenin, and DPD (**Fig 3B-F**). Individual CF ciliated cells isolated directly from turbinate brushings also exhibited less NO production compared with non-CF cells in response to multi-T2R agonist quinine (individual cell traces shown in **Fig 3G** and average of independent experiments shown in **Fig 3H**), suggesting (along with qPCR data above) that this is not due to reduced numbers of ciliated cells resulting in reduced T2R expression.

**Figure 3.**
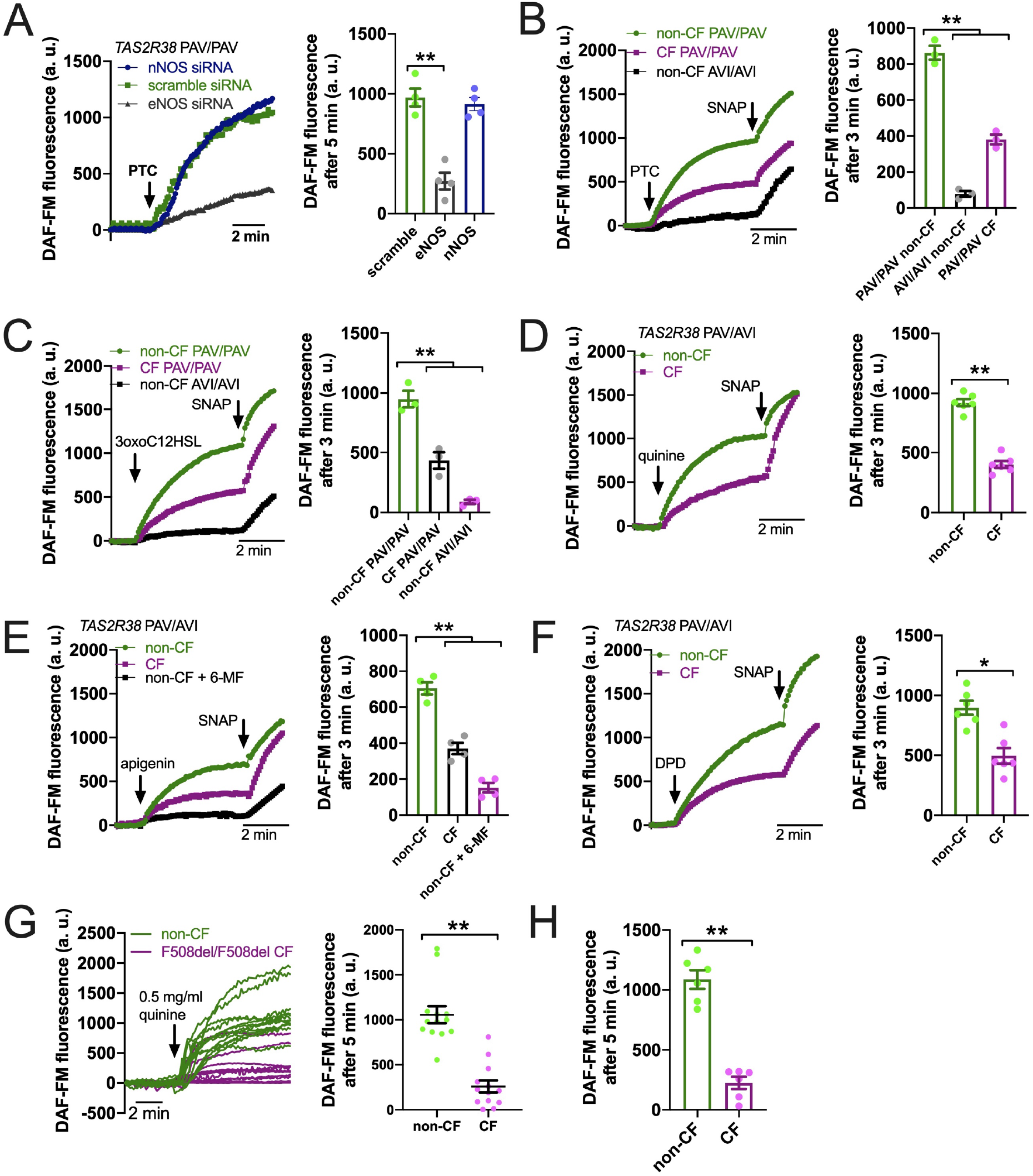
Reduced T2R-stimulated NO responses in CF nasal cells. **(A)** Left, representative traces of DAF-FM fluorescence, a dye that terminally reacts with NO and reactive nitrogen species and becomes fluorescent in primary nasal air-liquid interface cultures (ALIs). Nasal ALIs were genotyped as PAV/PAV (homozygous functional T2R38) and stimulated with T2R38 agonist phenylthiocarbamide (PTC). Cells were previously treated during differentiation with siRNA directed against endothelial or neuronal nitric oxide synthase (eNOS or nNOS, respectively) or scramble siRNA. Right shows bar graph of data from independent experiments using ALIs from separate individual patients. eNOS siRNA significantly reduced PTC-stimulated NO production by one-way ANOVA with Dunnett’s posttest (scramble siRNA as control). **(B)** Traces (left) and bar graph (right; each point represents an independent experiment from different patients) of DAF-FM fluorescence ALIs genotyped as *TAS2R38* PAV/PAV (homozygous functional) CF or non-CF as well as non-CF *TAS2R38* AVI/AVI (homozygous non-functional) during stimulation with T2R38 agonist PTC. Both non-CF AVI/AVI and CF PAV/PAV had reduced NO production in response to PTC vs non-CF PAV/PAV; **p<0.01 by one-way ANOVA with Bonferroni posttest. **(C)** Traces (left) and bar graph (right) similar to *B* but with cells stimulated with T2R38 agonist 3oxoC12HSL. Both non-CF AVI/AVI and CF PAV/PAV had reduced NO production vs non-CF PAV/PAV; **p<0.01 by one-way ANOVA with Bonferroni posttest. **(D)** Traces (left) and bar graph (right) showing DAF-FM responses in CF and non-CF ALIs genotyped as AVI/PAV *TAS2R38* during stimulation with multi-T2R agonist quinine. CF cells has reduced NO responses vs non-CF cells; **p<0.01 by Student’s *t* test. **(E)** Traces (left) and bar graph (right) showing DAF-FM responses in CF and non-CF ALIs genotyped as AVI/PAV *TAS2R38* during stimulation with T2R14 agonist apigenin. Both CF cells and non-CF cells pre-treated with 6-methoxyflavanone (6-MF) had reduced NO production vs non-CF cells; **p<0.01 by one way ANOVA with Bonferroni posttest. **(F)** Traces and bar graph similar to *D* but with T2R14 agonist diphenhydramine. Significance by Student’s *t* test; \**p*<0.05 **(G)** Left, representative traces from individual isolated ciliated cells from CF (magenta) and non-CF (green) middle turbinate. Right, plot of individual cells from representative experiments on left; **p<0.01 by Student’s t test. **(H)** Bar graph of data from individual independent experiments as in *G* from 5 separate CF and 5 separate non-CF patients. CF patient cells had reduced NO production; **p<0.05 by Student’s *t* test. In several panels, non-specific NO donor S-nitroso-N-acetyl-D,L-penicillamine (SNAP; 20 µM) was used as a positive control at the end of the experiment.

We questioned if the defect in NO production was a general defect in NO production, and tested this using small molecule Akt activator SC79, which results in phosphorylation and activation of eNOS in airway epithelial cells ((84) and **Supplemental Fig 3A**). SC79 stimulated NO production that was reduced by Akt inhibitors MK2206 and GSK690693 (**Supplemental Fig 3B**) as well as NOS inhibitor L-NAME (**Supplemental Fig 3B**). There was no difference in SC79-activated NO production with T2R38 genotype (**Supplemental Fig 3C**), and Akt inhibitors did not decrease NO during stimulation with T2R agonist 3oxoC12HSL (**Supplemental Fig 3D**). SC79 and 3oxoC12HSL activated NO production with different kinetics, with SC79 inducing a more sustained NO production and 3oxoC12HSL inducing more of a “burst” of higher level NO production (**Supplemental Fig 3E**), supporting that these two compounds activate eNOS through different mechanisms. We saw no difference in SC79 stimulated NO production between non-CF and F508del/F508del CF cells (**Supplemental Fig 3F**). This supports that there is not a general defect in eNOS function in CF cells, but rather there is a more specific defect in T2R signaling to eNOS downstream of T2R activation.

To confirm that this was really due to CFTR mutations and not another cause like a genetic linkage artifact, we tested NO production in 16HBE cells that were CRISPR modified to contain CFTR mutations, either F508del or premature stop codon G542X (66). 16HBE cells are an SV40 immortalized non-CF bronchial cell line (85) that produces NO in response to bitter taste agonist denatonium benzoate (86). CFTR has two polymorphisms at residue 470, M or V, that can affect protein stability and disease severity. The F508del mutation is almost exclusively paired with the M470 polymorphism (66). However, the Wt parental 16HBE cells contain V470 CFTR. We saw that both F508del M470 and F508del V470 CFTR 16HBE cells grown at ALI had reduced NO during stimulation with denatonium benzoate compared with the parent non-CF cells (**Supplemental Figure 4A-B**). Because of this, we used the more clinically relevant F508del M470 cell line for the rest of the studies here. As with F508del, G542X cells exhibited less NO production during denatonium stimulation compared with the parent non-CF cells (**Fig 4A**). Sodium benzoate was used as an osmotic/pH control. We ensured that DAF-FM traces reflected NOS activity using NOS inhibitor L-NAME and inactive control D-NAME (1 hr. pretreatment; 10 µM; **Fig 4A**). These data suggest that CFTR mutations alone can reduce NO signaling during T2R stimulation, even within an isogenic background.

**Figure 4.**
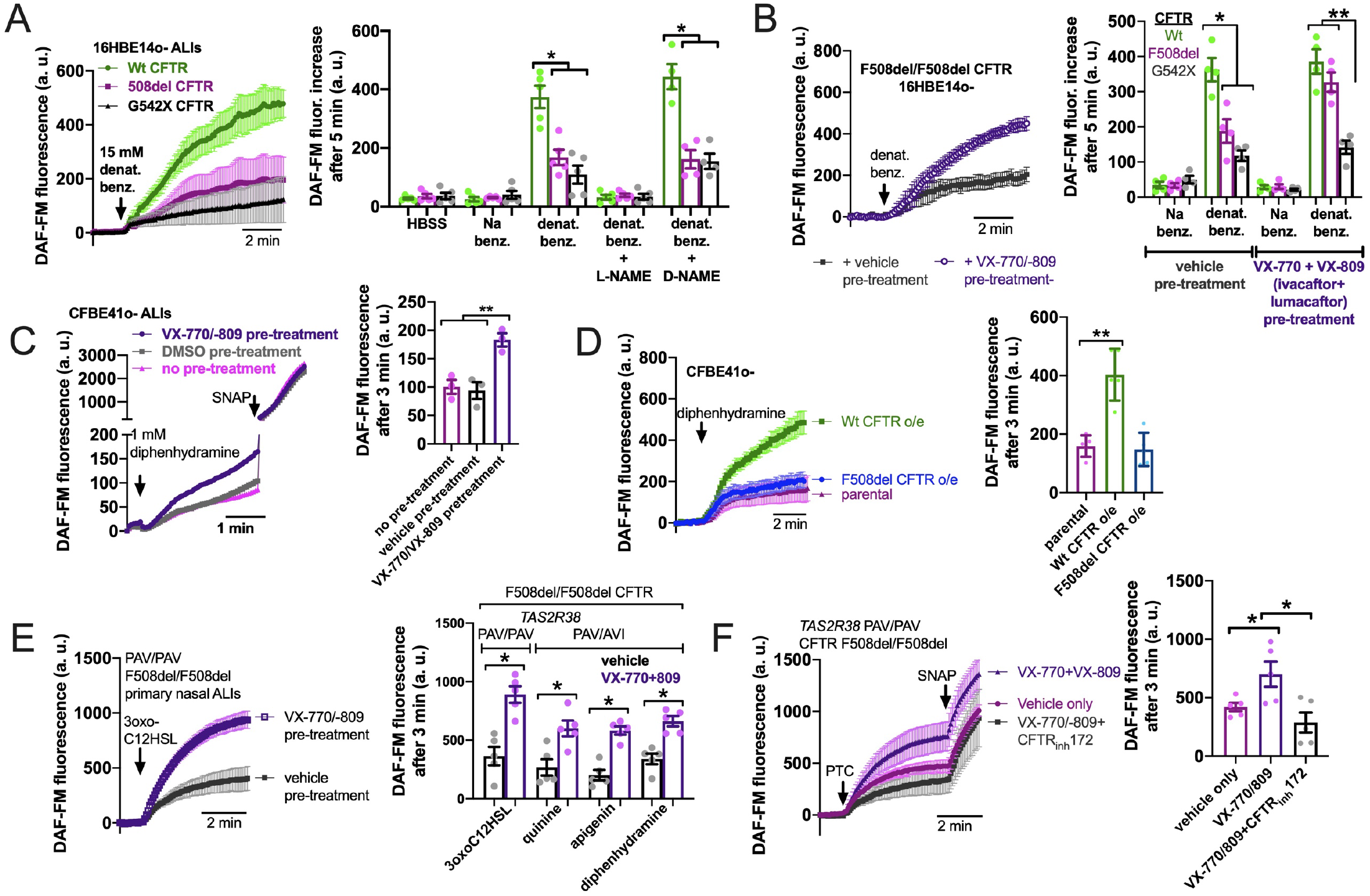
Loss of CFTR function reduces T2R-stimulated NO production in isogenic cell lines and restoration in cell lines and primary cells by CFTR corrector/potentiator treatment. **(A)** Left, DAF-FM traces of nitric oxide (NO) production in response to multi-T2R agonist denatonium benzoate in 16HBE14o-air liquid interface cultures (ALIs) with Wt CFTR (parental line) or clones modified by CRISPR/Cas9 to have homozygous F508del/F508del or G542X/G542X CFTR. Right, bar graphs of DAF-FM fluorescence changes in the three cell lines with T2R agonists denatonium ± NO synthase (NOS) inhibitor L-NAME or inactive analogue D-NAME. Equimolar sodium benzoate is used as control for pH and osmolarity. CFTR modified cells exhibited reduced NO production by one-way ANOVA with Bonferroni posttest; *p<0.05. NO production was blocked by L-NAME but not D-NAME. Each data point represents one independent experiment using cells from a different passage. **(B)** Left, traces of denatonium-stimulated NO production (DAF-FM) after VX-770+VX-809 or vehicle (0.2% DMSO) pre-treatment. Right, bar graph showing changes in DAF-FM fluorescence in cells with Wt CFTR, F508del CFTR, or G542X CFTR after VX-770+VX-809 or vehicle pre-treatment. Significance determined by one-way ANOVA with Bonferroni posttest; \**p*<0.05 and \*\**p*<0.01. Each data point represents one independent experiment using cells from a different passage. **(C)** CFBE41o-cell (homozygous F508del parental line) ALIs exhibited increased NO production in response to T2R14 agonist diphenhydramine (DPD) after treatment with VX-770/VX-809. Left, representative traces. Right, bar graph showing data points from independent experiments. Significance by one-way ANOVA with Bonferroni posttest; \*\**p*<0.01. **(D)** Overexpression of Wt CFTR but not F508del CFTR increased NO production in CFBE41o-cells. Left, representative traces. Right, bar graph showing data points from independent experiments using cells from different passages. Significance by one-way ANOVA with Bonferroni posttest; \*\**p*<0.01. **(E)** Left, trace showing DAF-FM in primary nasal ALIs from F508del/F508del CF *TAS2R38* PAV/PAV patients, where VX-770+VX-809 increased NO production in response to T2R38 agonist 3oxoC12HSL. Right, bar graph showing data points from independent experiments using cells from different CF patients. PAV/AVI cells were used to test multi-T2R agonist quinine or T2R14 agonists apigenin and diphenhydramine after vehicle or VX-770+VX-809 treatment. Significance by one way ANOVA with Bonferroni posttest. **(F)** Left, traces of DAF-FM in primary nasal ALIs from F508del/F508del CF *TAS2R38* PAV/PAV patients stimulated with T2R38 agonist PTC after pre-treatment with vehicle only, VX-770+VX-809, or VX-770+VX-809+CFTR_inh_172. Right, bar graph showing data points of DAF-FM fluorescence increases from independent experiments using cells from different CF patients. Increased DAF-FM fluorescence changes with VX-770+VX-809 were reduced with CFTR_inh_172 by one-way ANOVA with Bonferroni posttest; *p<0.05. In several panels of this figure, non-specific NO donor S-nitroso-N-acetyl-D,L-penicillamine (SNAP; 20 µM) was used as a positive control at the end of the experiment.

To further test if CFTR function is required for T2R to NO signaling, we treated F508del CFTR 16HBE ALI cultures for 48 hrs with corrector/potentiator combination VX-770 (ivacaftor; 1 µM) and VX-809 (lumacaftor; 3 µM) or vehicle (DMSO) alone (**Fig 4B**). VX-770 + VX-809 pre-treatment increased NO production during denatonium stimulation (**Fig 4B**). Wt and G542X cells were unaffected. Corrector/potentiator therapies cannot restore CFTR activity with premature stop codon mutations, so the lack of effect on G542X cells suggests that the effects of VX-770 + VX-809 is dependent on their ability to restore CFTR function. Using a Cl-substitution SPQ assay (69) (**Supplemental Fig 5A**), we confirmed that VX-770 + VX-809 treatment restored substantial apical membrane cAMP-stimulated ion permeability in F508del but not G542X cells (**Supplemental Fig 5B**). We also found that VX-770 + VX-809 did not affect T2R Ca^2+^ responses in F508del 16HBE cells (**Supplemental Fig 6**), ruling out increased Ca^2+^ as a mechanism for the effects of VX-770 + VX-809 on NO production.

**Figure 5.**
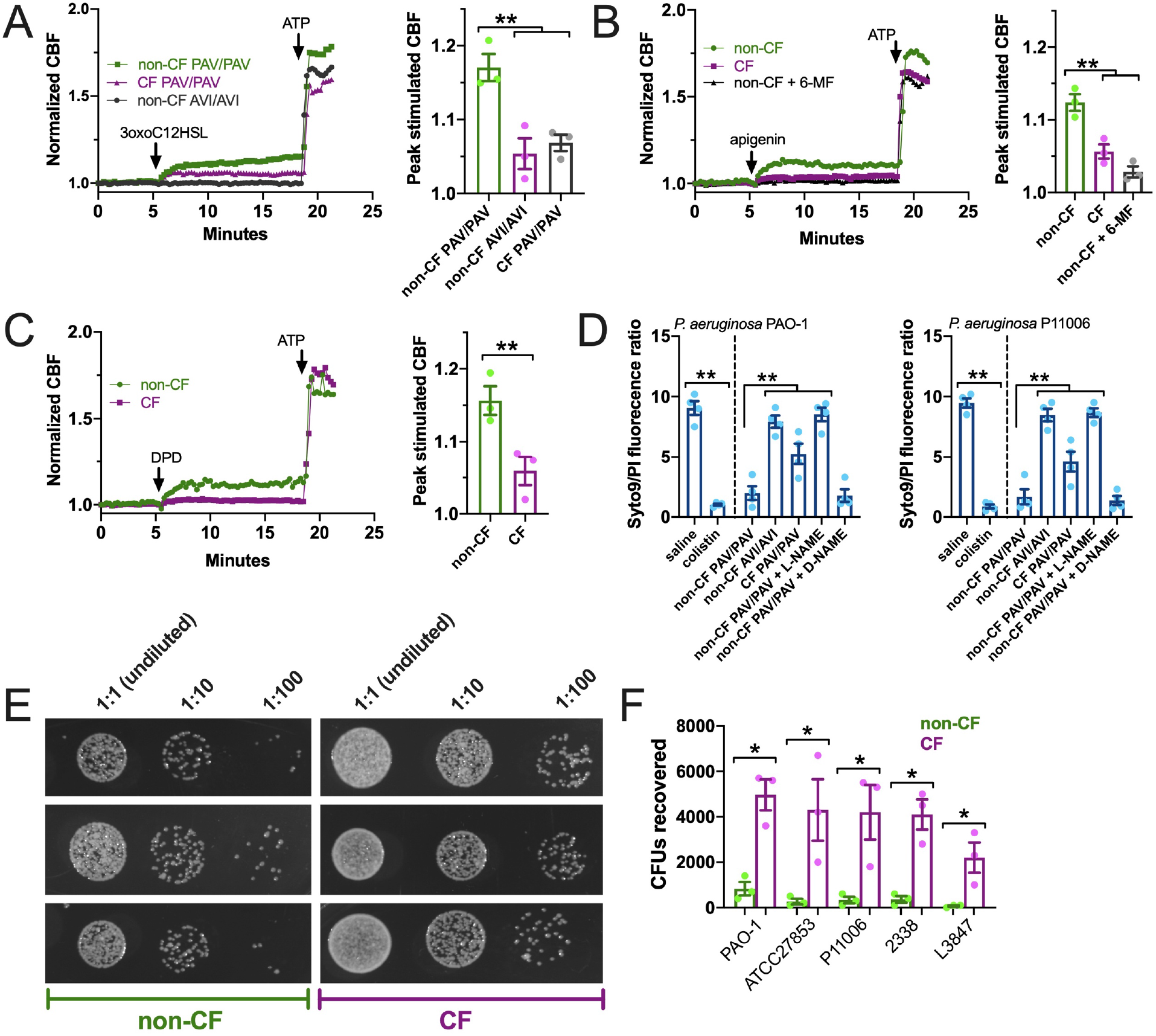
T2R ciliary and antibacterial responses are reduced in CF patient ALIs. **(A-C)** Whole-field ciliary beating was quantified during stimulation with T2R38 agonist 3oxoC12HSL, apigenin, and diphenhydramine (DPD). Representative traces (left) and bar graphs showing independent experiments using ALIs from separate patients (right; n = 3) are shown. CF cells exhibited reduced ciliary responses vs non-CF cells; *p*<0.01 by one-way ANOVA with Bonferroni posttest (*A,B*) or Student’s t test (*C*). ALIs in *A* were genotyped as *TAS2R38* PAV/PAV (functional) or AVI/AVI (non-functional). ALIs in *B* and *C* were from PAV/AVI patients. T2R14 agonist 6-methoxyflavanone was used as a control for apigenin as described earlier in the text. **(D)** Bacteria Syto9 (live) and propidium iodide (dead) staining were quantified following bacterial killing assays as described in the text. Saline and antibiotic colistin were used as negative and positive controls, respectively. Significance determined by one-way ANOVA with Bonferroni posttest; **p<0.01. CF cells killed less bacteria (higher Syto9/PI ratio) than non-CF cells. Non-CF cell killing was reduced by L-NAME but not inactive D-NAME. Both lab PAO-1 and clinical P11006 strains were used. **(E)** Images of PAO-1 CFUs recovered from three CF and non-CF ALIs from individual patients. **(F)** Quantification of independent experiments (n=5) as in *E* using five strains of *P. aeruginosa* showing reduced bacterial killing from CF cells. Significance by one-way ANOVA with Bonferroni posttest; \**p*<0.05.

**Figure 6.**
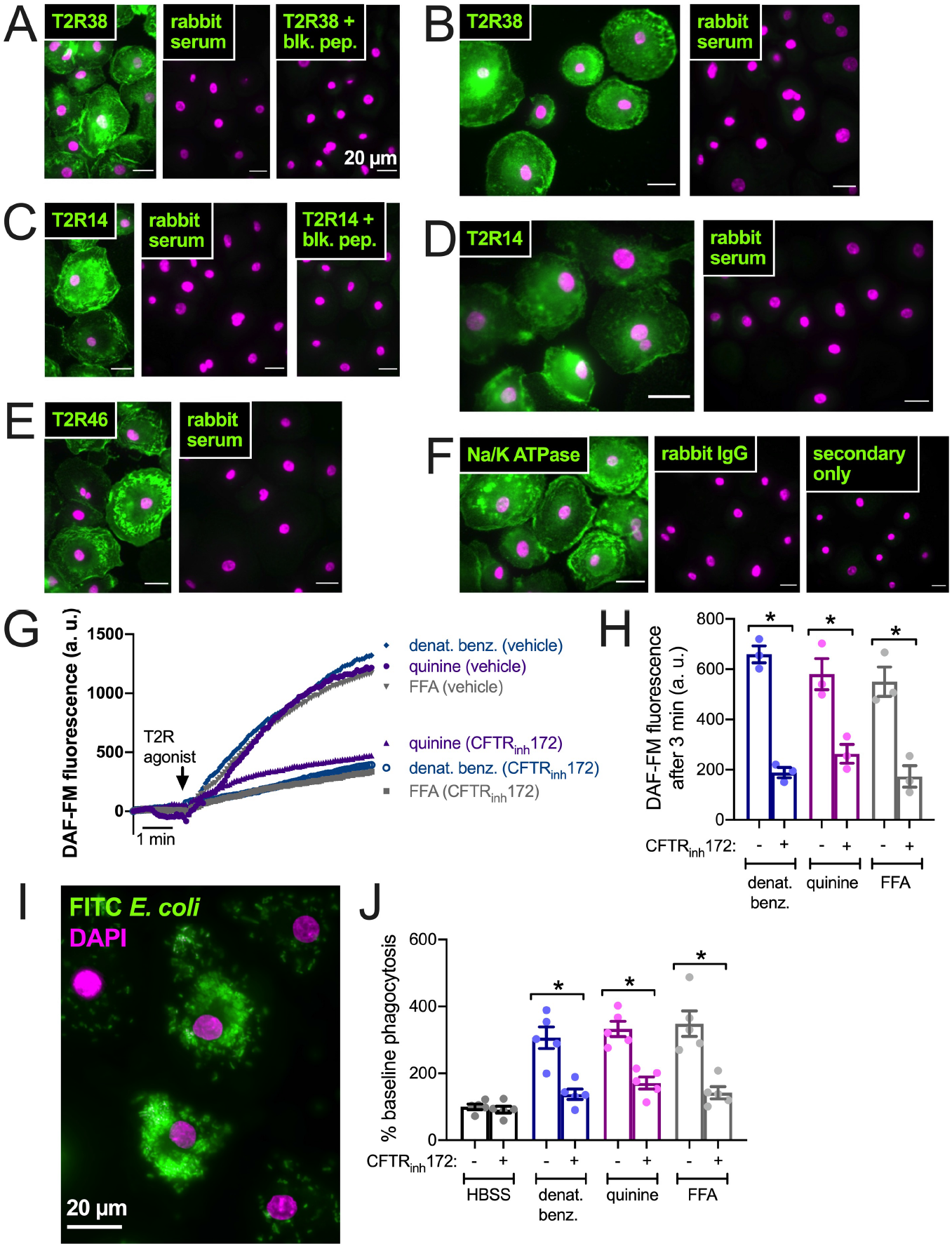
CFTR inhibition results in reduced T2R-stimulated NO production and NO/cGMP phagocytosis in human macrophages. **(A)** Macrophages exhibited staining reminiscent of plasma membrane ruffles with C-terminal T2R38 antibody (left panel) that was blocked in the presence of T2R38 blocking peptide (right panel). **(B)** A different T2R38 antibody resulted in similar staining. **(C)** Macrophages exhibited staining reminiscent of plasma membrane ruffles with C-terminal T2R14 antibody (left panel) that was blocked in the presence of T2R14 blocking peptide (right panel). **(D-F)** A different T2R14 antibody (*D*) and T2R46 antibody (*E*) also resulted in similar staining reminiscent of Na^+^/K^+^ ATPase plasma membrane staining (*F*). All data in *A-F* are representative of images taken from macrophages from >3 different donors stained in 3 different experiments. Affinity purified rabbit serum was used as a negative control for non-specific staining of all polyclonal antibodies. **(G-H)** NO production was measured using DAF-FM. Cells stimulated with denatonium benzoate, quinine, or flufenamic acid (FFA) exhibited reduced NO production after incubation with CFTR_inh_172. Representative traces shown in *G* and bar graph in *H* shows results from independent experiments. **(I)** Representative image of macrophage phagocytosis of FITC-labeled E. coli. **(J)** Bar graph of phagocytosis assay quantification via plate reader, showing increase in phagocytosis in macrophages stimulated with multi-T2R agonists denatonium benzoate or quinine as well as T2R14 agonist FFA, all *p*<0.01 vs HBSS buffer only control by one-way ANOVA with Bonferroni posttest. Increased phagocytosis was reduced after pre-incubation with CFTR_inh_172; *p<0.05 by one-way ANOVA/Bonferroni posttest.

To further confirm effects of corrector potentiator therapies, we tested CFBE41o-cells, also an SV40 immortalized bronchial cells but from a F508del/F508del CF patient lung (63). CFBE41o-cells produce NO in response to T2R14 agonist DPD (72). This NO production was increased after VX-770 + VX-809 treatment (**Fig 4C**). Even further supporting the role of CFTR in T2R NO production, we tested CFBE41o-cells stably over-expressing Wt or F508del CFTR. Wt CFTR expression increased DPD-stimulated NO production while F508del was without effect (**Fig 4D**). Lastly, F508del/F508del primary CF cells pretreated with VX-770 + VX-809 exhibited increased NO production to T2R agonists 3oxoC12HSL, quinine, apigenin, and DPD (**Fig 5E**). Interestingly, co-application (48 hrs) of VX-770 + VX-809 and CFTR_inh_172 (5 µM) abrogated the effect of VX-770 + VX-809 (**Fig 4F**).

How does this affect innate immunity in CF cells? To answer this, we first tested ciliary beat frequency (CBF), which drives mucociliary transport and clearance of inhaled/inspired pathogens. T2R activation and stimulation of NO production activates guanylyl cyclase to increase CBF via protein kinase G (22, 74, 87). CF ALIs exhibited reduced CBF increases with 3oxoC12HSL (**Fig 5A**), apigenin (**Fig 5B**), and DPD (**Fig 5C**) compared with non-CF ALIs. Because the whole field analysis of CBF takes into account only actively beating areas as determined by the software (77), this further confirms that effects observed here are not due to differences in numbers of ciliated cells, but rather defects intrinsic to the ciliated cells themselves.

Next, we examined the ability of CF and non-CF nasal ALIs to kill both lab (strain PAO1) and clinical (strain P11006) *P. aeruginosa*. In this assay, incubation of *P. aeruginosa* with nasal ALIs results in bacterial killing over 2 hours that requires T2R38 and NO production, as previously demonstrated (39, 74). PAV/PAV (functional T2R38) non-CF cultures killed both strains of *P. aeruginosa* to levels comparable with antibiotic colistin (**Fig 5D**). AVI/AVI non-CF cultures and PAV/PAV CF cultures both exhibited reduced bacterial killing (**Fig 5D**). NOS inhibitor L-NAME, but not inactive D-NAME, inhibited non-CF nasal epithelial cell bacterial killing (**Fig 5D**), supporting the role for NO in this assay. Together, these data suggest that CF nasal epithelial cells have a reduced capacity to both clear and kill bacteria during T2R stimulation.

We next tested if this was a general effect of CFTR function on T2R signaling or if this was specific to nasal epithelial cells. We previously reported that macrophages exhibit T2R-induced Ca^2+^ signaling and NO production downstream of eNOS (42, 74). Instead of regulating ciliary beating, this pathway instead regulates phagocytosis. We confirmed that macrophages express T2R38 (**Fig 6A-B**), T2R14 (**Fig 6C-D**), and T2R46 (**Fig 6E**) with localization reminiscent of plasma membrane marker Na^+^/K^+^ ATPase (**Fig 6F**). We examined NO production by DAF-FM imaging and found that pre-treatment (24 hrs) with CFTR_inh_172 (10 µM) resulted in reduced NO production when stimulated with multi-T2R agonists denatonium benzoate or quinine as well as T2R14-specific agonist flufenamic acid (FFA; **Fig 6G-H**). This was accompanied by reduced phagocytosis of FITC-labeled *E. coli* (**Fig 6I-J**), suggesting that loss of CFTR function can reduce T2R signaling in multiple cell types.

## Discussion

We show above that T2R-mediated NO generation is reduced in cells with compromised CFTR function. The mechanism underlying this defect is not yet known but appears to be downstream of Ca^2+^, as T2R-induced Ca^2+^ responses are normal. This is also not likely due to reduced Akt signaling with loss of CFTR function, as was previously suggested for the reduced eNOS activation in CF endothelial cells downstream of sheer stress (35). T2R-dependent NO production was Akt-independent, and NO production was not different between CF and non-CF cells when these cells were stimulated with small-molecule Akt activator SC79 (84, 88). Further studies are needed to identify altered pathway component(s) of T2R signaling to eNOS in CF cells. Regardless, this suggests an intrinsic reduction in CF cell innate immunity via reduced T2R NO signaling caused by loss of CFTR function. Further investigation into the downstream signaling of T2Rs to eNOS is required to understand the mechanism(s) of the alteration(s) observed in CF cells. Additionally, a caveat to our study is that the patient numbers observed here are limited by the need to genotype and segregate CF patients for T2R38 status. While large for an *in vitro* airway cell biology study, the numbers here are small compared with clinical studies and results will require confirmation in larger patient cell sets.

Notably, while NO is involved in T2R signaling in airway epithelial cells and macrophages, it does not appear to be required for taste signal transduction in taste buds (89). T2R-stimulated Ca^2+^ signals in the type II taste cell activate membrane depolarization to cause release of ATP through CAHLM1/3 ion channels to activate purinergic receptors on afferent gustatory neurons (90). Patients with CF have been suggested to have an increased taste sensitivity (91), but this is across all taste modalities (sweet, bitter, salty, sour) with different chemosensory mechanisms. Another study determined CF patient taste was not different from non-CF individuals (92). Regardless, because we find a difference downstream of the actual T2R receptor Ca^2+^ response at the level of NO signaling, the lack of a reduction of bitter taste perception does not discount or contradict our observations here.

While it is certainly possible that effects observed are due to different inflammatory environments of CRS vs CF-CRS from which the cells were removed, we do not believe this is the case. We observe minimal differences in phenotypes of CRS vs non-CRS cells after ∼4-6 weeks of culturing and differentiation. Phenotype is overwhelmingly based on genetics after culturing in the same media for several weeks. As all cells used here were from tissue that had been propagated in the same media, we would not expect any significant lasting effects of the *in vivo* inflammatory patient environment independent of patient genetics. We also demonstrate here that CFTR corrector/potentiator treatment *in vitro* can restore T2R NO production to non-CF cell levels. This suggests that the T2R innate immune arm is enhanced in patients receiving these small molecule therapies. This also supports that the defect in NO production stems from defective CFTR function and not from inflammatory differences.

However, T2R-induced NO production is not restored in patients who cannot benefit from small molecule therapies, such as those with G542X CFTR mutations. We hypothesize that enhancement of NO production via another method (e.g., targeting the Akt pathway as with SC79 or artificial NO donor compounds), may be beneficial to reducing respiratory infections with gram-negative bacteria, though further research is needed to test this in more detail. Notably, while ion permeability was increased by corrector/potentiator treatment, it remained much lower than non-CF cells (e.g., **Supplemental Figure 5C**), as expected from other studies in these cells (66). Nonetheless, NO levels were restored to near the levels of non-CF cells (e.g., **Fig 4B**). This result suggests some level of CFTR function is required for T2R NO production but not necessarily levels comparable to non-CF cells. We speculate that this may relate to CFTR’s proposed function as a scaffolding protein for signaling (19). While CFTR_inh_172 reduced T2R-mediated NO production after VX-770 + VX-809 treatment, CFTR_inh_172 can lead removal of CFTR from the plasma membrane and/or CFTR degradation (93). It remains to be determined if Cl^−^ conductance or changes in intracellular [Cl^−^] specifically affect T2R NO production.

The reduction of T2R-mediated NO generation upon loss of CFTR function may explain discrepancies and/or subtle effects of *TAS2R* genetics on CF disease progression. This may also contribute to the specific susceptibility of CF patients to gram-negative bacteria like *P. aeruginosa*, which (1) secrete the AHL and quinolone agonists known to activate these receptors and (2) are most sensitive to bactericidal effects of nasal epithelial cell generated NO production. We also hypothesize that restoration of CFTR function by small molecule corrector/potentiator therapies, which restore T2R-mediated NO production in response to bacterial ligands, may enhance the importance of *TAS2R* polymorphisms in CF patient respiratory infections. Future studies of *TAS2R* polymorphisms and outcomes in CF patients receiving such therapies are needed to understand if T2Rs are more involved in respiratory defense in these patients. Additional work may reveal that patients with reduced T2R38 function can benefit from topical application (e.g., nasal lavage) of agonists for other T2Rs like T2R14 (72).

## Acknowledgements

This study was supported by a Cystic Fibrosis Foundation grant (LEE21G0 to R.J.L.) and NIH grants R01DC016308 (to R.J.L.) and R01AI167971 (to N.D.A., J.N.P., and R.J.L.). We thank M. Victoria, B. Hariri, and J. Freund (University of Pennsylvania Perelman School of Medicine) for technical assistance with cell culture. We thank B. Chen, N. Cohen, and L.E. Kuek (University of Pennsylvania Perelman School of Medicine) for assistance with isolation of patient nasal cells. We thank J. Riley (University of Pennsylvania Perelman School of Medicine, Human Immunology Core, supported by P30CA016520 and P30AI045008) for access to primary human monocytes. We thank H. Valley and M. Mense (Cystic Fibrosis Foundation, CFFT Lab) for 16HBE CRISPR-modified cell lines.

