## Supplemental Material for "Bitter taste receptor-stimulated nitric oxide innate immune responses are reduced by loss of CFTR function in nasal epithelial cells and macrophages"

### Supplemental Figure 1

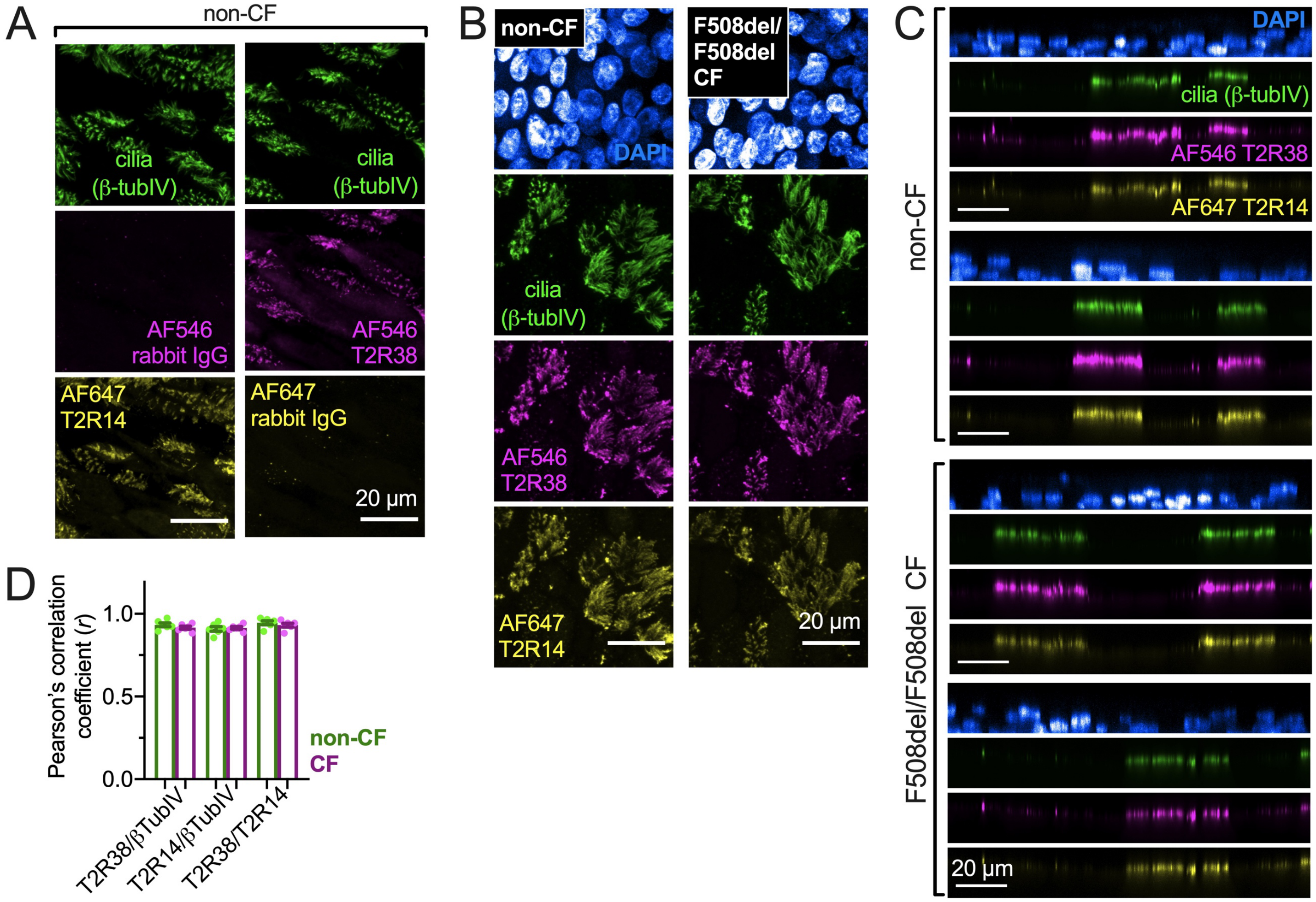


**Supplemental Figure 1. Immunofluorescence localization of T2R38 and T2R14 in CF and non-CF nasal cilia. (A)** Representative images showing staining of two non-CF patient ALIs with AlexaFluor (AF) 546 labeled T2R38 primary antibody, AF647 labeled T2R14 primary antibody, or AF546 or AF647 labeled rabbit serum control. Representative of results from 3 independent experiments using cells from 3 non-CF patients. **(B)** Representative images comparing staining of AF546-labeled T2R38 antibody and AF647-labeled T2R14 antibody in non-CF (left) and CF (right) patient ALIs. **(C)** Orthogonal plans from images as in *C*, showing similar localization in CF and non-CF cells. Images taken at 60x 1.4NA objective with 0.2 μm *z* step size, not corrected for refractive index mismatch to minimize image processing. Images are representative of 3 independent experiments using cells from 3 CF and 3 non-CF patients. **(D)** Pearson's correlation coefficient was computed using ImageJ showing strong correlation ($r \geq 0.9$) of the T2R14 and T2R38 signals as well as either T2R14 and T2R38 signals with β-tubulin IV. No significant differences were observed between CF and non-CF patients by ANOVA. All results shown are representative of 4 ALIs from 4 individual CF and 4 individual non-CF patients.

### Supplemental Figure 2

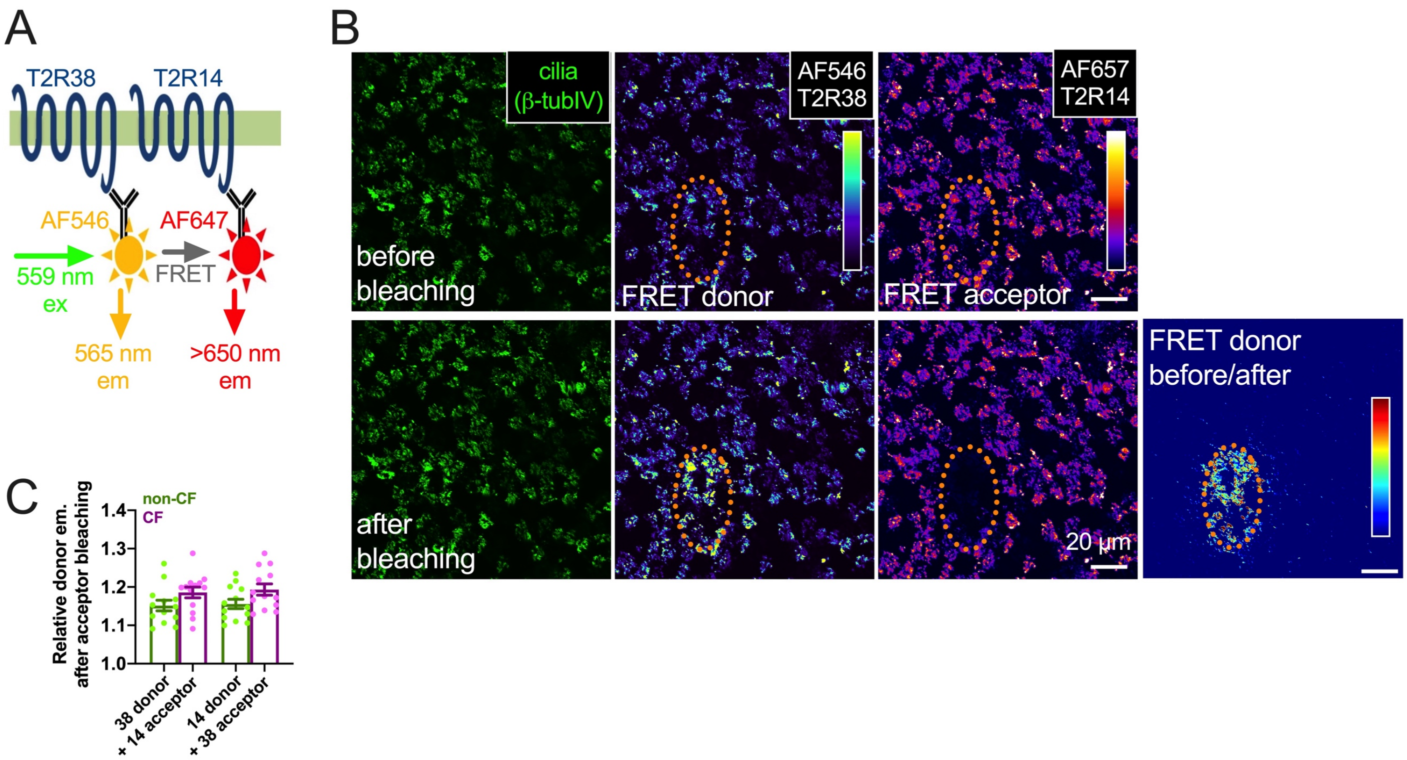


**Supplemental Figure 2. Confirmation of similar co-localization of T2R14 and T2R38 by quantification of antibody-based Förster resonance energy transfer (FRET). (A)** Co-localization was quantified by measurement of FRET efficiency of labeled primary antibodies as described in (1, 2). **(B)** Bleaching of a region of the FRET acceptor via 647 laser and tornado bleaching function in Olympus Fluoview (photo-bleached area shown in yellow circle) resulted in an increase in fluorescence of the donor, confirming FRET. **(C)** Quantification of FRET efficiency (the relative increase in donor fluorescence after acceptor bleaching; E). as described in the methods revealed similar FRET between T2R38 and T2R14 CF and non-CF cells. Results representative of 12 independent experiments using ALIs grown from 4 CF and 4 non-CF patients (3 ALIs each). E was not significantly different between the two genotypes. As described (1), FRET efficiency decreases to the 6$^{th}$ power as distance (R) increases, thus $R = R_o \bullet [(1\text{-}E)/E]^{1/6}$ where $R_o$ is the distance at which the E is 50% (51 Å for AF555 and AF647). A ~0.16 E equates to ~67 Å in distance between the two fluorophores. Together with Figure 2 in the main text, these data suggest no major changes in T2R localization between CF and non-CF airway epithelial cells.

### Supplemental Figure 3

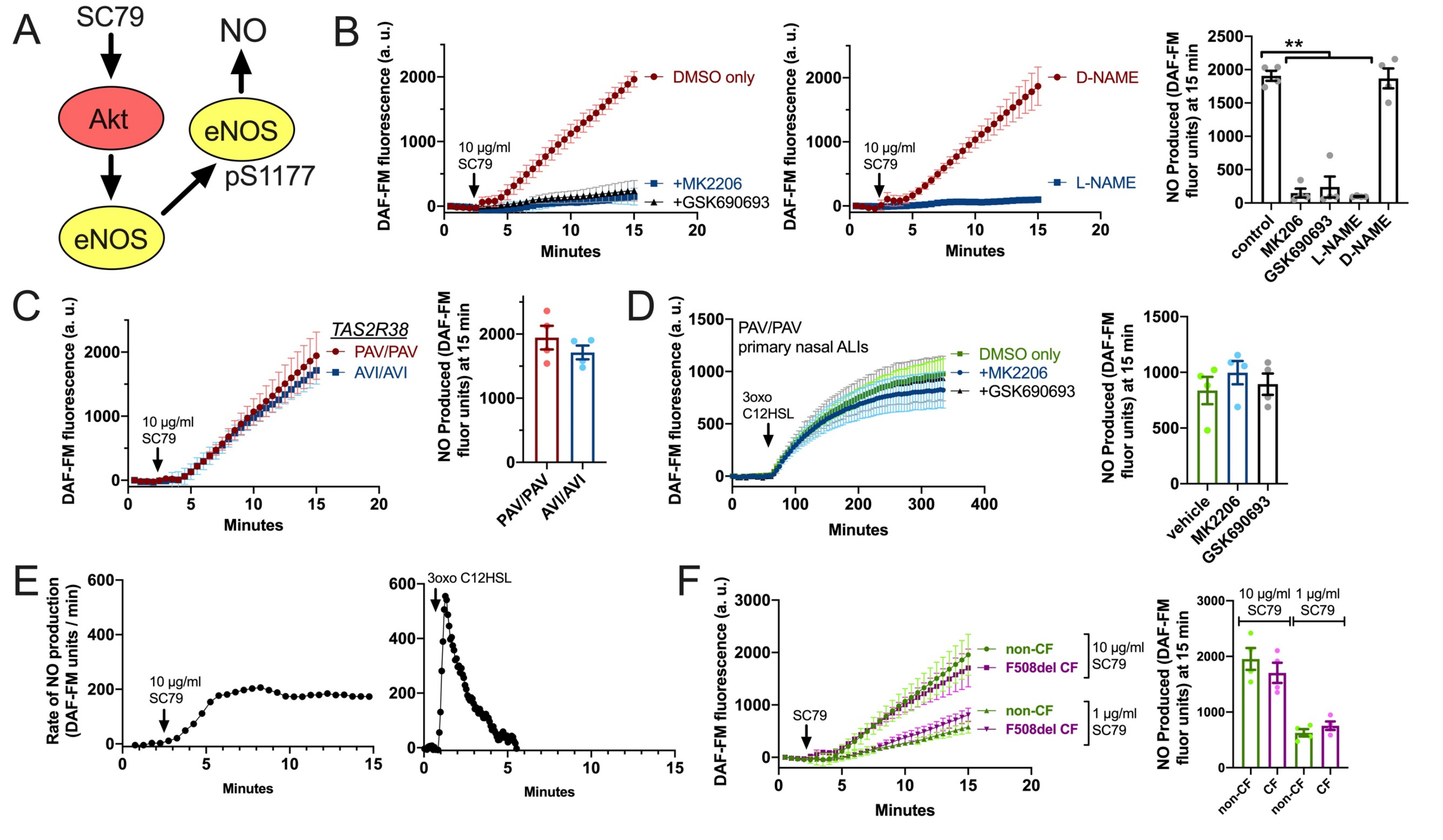


**Supplemental Figure 3. Intact Akt-stimulated NO production in CF nasal epithelial cells. (A)** It was previously demonstrated (3) that Akt activator SC-79 causes eNOS phosphorylation at S1177 in airway epithelial cells, stimulating NO production. **(B)** Left and middle, traces of DAF-FM fluorescence increases in response to 10 µg/ml SC79 showing increased in NO production that is inhibited by Akt inhibitors MK2206 and GSK690693 (left; 1 µM each for 45 min pre-treatment) or NOS inhibitor L-NAME (right; 10 µM for 45 min pre-treatment) but not inactive analogue D-NAME. Right is bar graph showing data from independent experiments using cells from different non-CF individuals. Significance by one-way ANOVA with Dunnett’s posttest comparing all values to control; $^{**}p<0.01$ **(C)** SC79/Akt-stimulated NO production is not different in cells genotyped for *TAS2R38* PAV/PAV or AVI/AVI, confirming that the SC79 response is independent of T2R38 function. No significant difference by Student’s *t* test. **(D)** Trace (left) and bar graph of independent experiments using ALIs from different patients (right) of DAF-FM fluorescence in PAV/PAV *TAS2R38* genotyped cells with 3oxoC12HSL (T2R38 agonist; 100 µM). No inhibition was observed with Akt inhibitors MK2206 or GSK690693, confirming T2R38 activation of eNOS is independent of Akt. No significant difference by one way ANOVA. **(E)** Forward derivative of representative DAF-FM fluorescence (showing rate of DAF-FM fluorescence change, roughly equating relative rates of NO production) during stimulation with SC79 or 3oxoC12HSL (100 µM) showing different kinetics of rate of NO production. SC79 activated more sustained NO production while T2R stimulation activated more of a burst of NO production that tapered off. **(F)** Representative traces (left) and bar graph of independent experiments using cells from different patients (right) showing no difference between CF and non-CF cells in terms of NO production with two different concentrations of SC79. No significant differences by one-way ANOVA. Together, these data suggest that Akt-activated NO production is intact in CF cells, and suggest there is a specific defect in T2R-activated NO production rather than a general defect in the ability of CF cells to produce NO. These data also support that the DAF-FM dye itself is not exhibiting different sensitivity in CF vs non-CF cells underlying the changes observed in the main text.

### Supplemental Figure 4

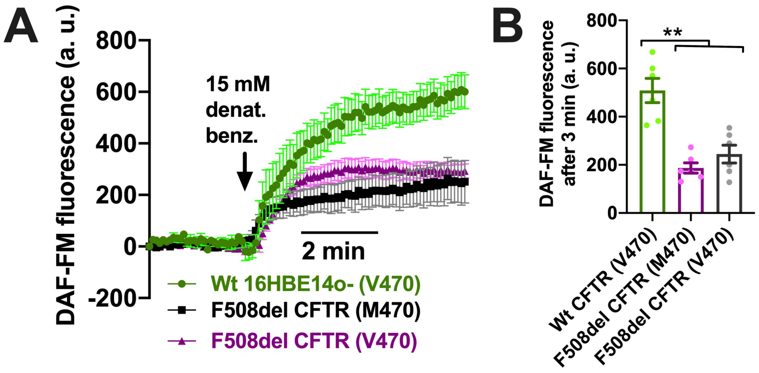


**Supplemental Figure 4. DAF-FM fluorescence changes were reduced in both F508del M470 and V470 cells. (A)** DAF-FM fluorescence traces showing increases in responses to multi-T2R agonist denatonium benzoate in cells with Wt CFTR, F508del M470, or F508del V470, generated as described (4). **(B)** Bar graph showing data from independent experiments (n = 5) showing reduced DAF-FM fluorescence change in both F508del M470 and V470, which were not significantly different from each other. Significance by one way ANOVA with Bonferroni posttest; ***p*<0.01.

Supplemental material for Carey, *et al.*, “Bitter taste receptor-stimulated nitric oxide innate immune responses are reduced by loss of CFTR function in nasal epithelial cells and macrophages.”

### Supplemental Figure 5

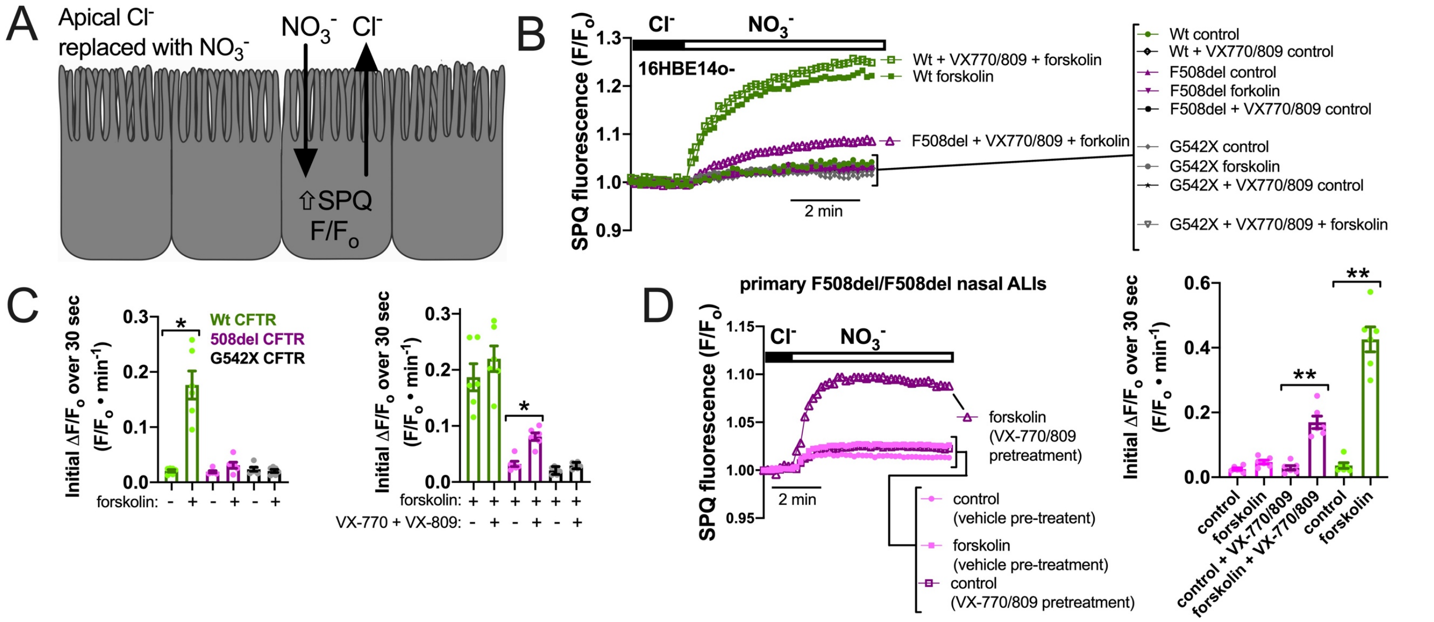

**Supplemental Figure 5: Increased apical $Cl^-$ permeability in CF cells treated with VX-770+VX-809. (A)** Diagram of NO substitution assay (used in (5)). SPQ is quenched by $Cl^-$ but not by $NO_3^-$. Most $Cl^-$ channels, including CFTR, have a nearly equal permeability to $Cl^-$ and $NO_3^-$. Apical substitution of $NO_3^-$ for $Cl^-$ results in electroneutral efflux of $Cl^-$ and influx of $NO_3^-$ along diffusional gradients, resulting in an increase in intracellular SPQ fluorescence (expressed as fluorescence normalized to fluorescence at time 0 [$F/F_o$]). **(B)** Representative traces of SPQ during apical $NO_3^-$ substitution ± forskolin stimulation, which resulted in an increase in the rate of SPQ fluorescence change (roughly equal to an increase in the relative apical anion permeability) in 16HBE14o- cells with Wt (parental strain), F508del (CRISPR modified), G542X (CRISPR modified) CFTR (4). **(C)** Bar graph from experiments as in *B* showing data points of initial rate of SPQ fluorescence change from 5-7 independent experiments. Significance by one-way ANOVA with Bonferroni posttest; *$p<0.05$. Non-CF (Wt CFTR parental 16HBEs) cells showed an increase in the rate of SPQ fluorescence change with forskolin stimulation. F508del and G542X CFTR cells did not. VX-770+VX-809 pre-treatment resulted in a significant increase in forskolin-stimulated SPQ change in F508del but not G542X cells. **(D)** Trace and bar graph showing experiments similar to *B* and *C* except using primary nasal ALIs from F508del/F508del CF patients. CF patients shown in pink. Bar graph also shows non-CF experiments shown in green, which exhibited an increase in SPQ fluorescence change with forskolin stimulation. F508del/F508del cells only showed a forskolin-stimulated increase after pre-treatment with VX-770+VX-809. These results confirm that VX-770+VX-809 increased apical anion permeability in F508del but not G542X cells, suggesting corrector/potentiator treatment increased F508del CFTR activity.

### Supplemental Figure 6

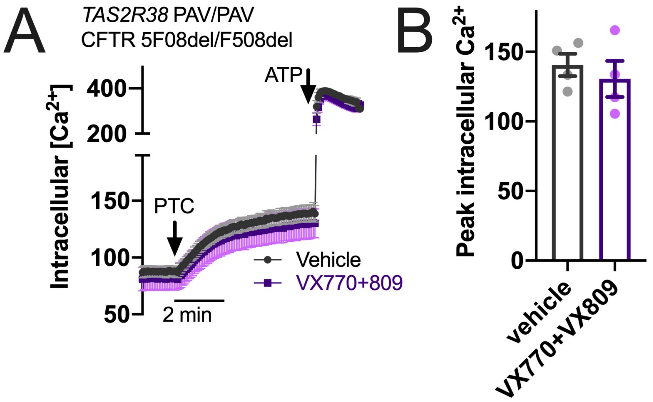


**Supplemental Figure 6. PTC-evoked $Ca^{2+}$ increases were not different in *TAS2R38* PAV/PAV CF ALIs after pre-treatment with V-X770+VX-809. (A)** Cells were loaded with Fura-2 identically as described in the text for Fluo-4 and as in (1). Trace showing intracellular $Ca^{2+}$ (calibrated from Fura-2 340/380 ratio as described in (1) using the method of Grynkiewicz (6)). Both resting $Ca^{2+}$ and stimulated $Ca^{2+}$ appeared the same. **(B)** Bar graph showing peak PTC-stimulated $Ca^{2+}$ from independent 4 experiments using ALIs from 4 different CF PAV/PAV patients. No significant difference by one-way ANOVA. These data suggest that VX-770+VX-809 pretreatment is not increasing T2R-stimulated NO production in CF cells due to enhancement of T2R $Ca^{2+}$ signaling.

### Supplemental Table 1

| Patient | Age at Surgery | Gender | Ethnicity | Diagnosis | # Prior FESS | Polyps | Lund-Mackay | SNOT-22 | Smoking History | Asthma | AFS | Abx | Steroids | Comorbidities | CFTR genotype |
|---|---|---|---|---|---|---|---|---|---|---|---|---|---|---|---|
| CF1 | 25 | Female | Caucasian | CF, CRS | 1 | No | 18 | 67 | No | Yes | No | Yes | Yes | CF, GERD, Asthma | F508del/1833delT |
| CF2 | 28 | Female | Caucasian | CF, CRS | 1 | No | 10 | 78 | No | Yes | No | Yes | Yes | CF, Asthma, GERD, DM | F508del/F508del |
| CF3 | 38 | Female | Caucasian | CF,CRS | 1 | Yes | 16 | 93 | No | No | No | No | Yes | CF, Lung transplant, DM | F508del/F508del |
| CF4 | 42 | Male | Caucasian | CF, CRS | 1 | No | 14 | 33 | No | Yes | No | Yes | No | CF, Lung transplant, DM, GERD, HTN | F508del/E585X |
| CF5 | 26 | Female | Caucasian | CF, CRS | 1 | No | N/A | 78 | No | Yes | No | No | No | CF, GERD, DM | F508del/F508del |
| CF6 | 23 | Male | Caucasian | CF, CRS | 1 | Yes | N/A | N/A | No | No | No | No | No | CF, GERD | F508del/F508del |
| CF7 | 33 | Female | Caucasian | CF, CRS | 1 | Yes | 13 | 8 | No | No | No | No | No | CF, Lung Transplant, Allergies, GERD, HTN, DM | F508del/F508del |
| CF8 | 32 | Female | Caucasian | CF, CRS | 1 | Yes | N/A | 77 | No | Yes | No | No | No | CF, GERD | F508del/F508del |
| CF9 | 58 | Female | Caucasian | CF, CRS | 2 | Yes | 18 | 27 | No | Yes | No | No | No | Allergies, GERD, HTN | F508del/F508del |
| CF10 | 27 | Female | Caucasian | CF, CRS | 0 | No | N/A | 59 | No | Yes | No | No | No | CF | F508del/G542X |
| CF11 | 32 | Male | Caucasian | CF, CRS | 2 | Yes | N/A | 43 | No | Yes | No | No | Yes | CF | F508del/F508del |
| CF12 | 38 | Male | Caucasian | CF, CRS | 1 | No | N/A | 28 | No | No | No | No | No | CF, Allergies | F508del/F508del |
| CF13 | 22 | Male | Caucasian | CF, CRS | 9 | Yes | 20 | 15 | No | No | No | No | Yes | CF, Allergies, GERD, DM | F508del/F508del |
| CF14 | 41 | Male | Caucasian | CF, CRS | 0 | Yes | 14 | 32 | No | Yes | No | Yes | No | CF, Allergies, DM | F508del/F508del |

| Patient | Age at Surgery | Gender | Ethnicity | Diagnosis | # Prior FESS | Polyps | Lund-Mackay[1,2] | SNOT-22[3] | Smoking History | Asthma | AFS | Abx | Steroids | Comorbidities | CFTR genotype |
|---|---|---|---|---|---|---|---|---|---|---|---|---|---|---|---|
| nonCF1 | 37 | Male | Caucasian | CRS | 1 | Yes | 18 | 68 | Yes | No | No | No | No | Allergies, Sinonasal Trauma | N/A |
| nonCF2 | 24 | Female | Caucasian | CRS | 0 | No | 13 | 69 | No | No | No | No | No | Allergies, GERD | N/A |
| nonCF3 | 19 | Male | Caucasian | CRS | 1 | Yes | N/A | 41 | No | Yes | No | No | No | Allergies | N/A |
| nonCF4 | 21 | Male | Caucasian | Orbital decompression | 0 | No | 0 | 0 | Yes | No | No | No | No | N/A | N/A |
| nonCF5 | 32 | Male | Caucasian | CRS | 1 | Yes | 10 | 42 | No | No | No | No | Yes | Allergies, GERD | N/A |
| nonCF6 | 34 | Female | Caucasian | CRS | 0 | Yes | 6 | 34 | No | Yes | No | No | No | Allergies | N/A |
| nonCF7 | 28 | Female | Caucasian | CRS | 1 | Yes | 26 | 27 | No | No | No | No | Yes | DM | N/A |
| nonCF8 | 43 | Male | Caucasian | CRS | 2 | Yes | 20 | 58 | No | No | Yes | Yes | Yes | AERD, Allergies, HTN | N/A |
| nonCF9 | 39 | Male | Caucasian | CRS | 1 | Yes | 16 | 59 | No | Yes | No | No | Yes | DM | N/A |
| nonCF10 | 44 | Male | Caucasian | CRS | 2 | Yes | 7 | 22 | No | No | No | Yes | No | N/A | N/A |
| nonCF11 | 44 | Female | Caucasian | CRS | 0 | Yes | 20 | 30 | No | Yes | No | No | No | HTN, Diabetes | N/A |
| nonCF12 | 44 | Male | Caucasian | CRS | 1 | No | 12 | 24 | No | No | No | No | No | HTN | N/A |
| nonCF13 | 28 | Male | Caucasian | Sinonasal tumor | 0 | No | 4 | 40 | Yes | No | No | No | No | N/A | N/A |
| nonCF14 | 32 | Male | Caucasian | CRS | 0 | No | 3 | 53 | No | No | No | No | No | N/A | N/A |
| nonCF15 | 28 | Male | Caucasian | CRS | 1 | No | 3 | N/A | No | No | No | No | No | Allergies | N/A |
| nonCF16 | 21 | Female | Caucasian | CRS | 0 | No | 3 | 14 | No | No | No | No | No | GERD | N/A |
| nonCF17 | 44 | Female | Caucasian | CRS | 0 | No | 4 | 40 | Yes | No | No | No | No | Allergies | N/A |
| nonCF18 | 42 | Female | African American | CRS | 1 | No | 1 | 89 | No | Yes | No | No | No | Asthma, Allergies, PE | N/A |
| nonCF19 | 35 | Female | Caucasian | CRS | 0 | No | 4 | N/A | No | No | No | No | No | N/A | N/A |
| nonCF20 | 23 | Female | Caucasian | CRS | 0 | Yes | N/A | 39 | No | Yes | No | No | No | N/A | N/A |
| nonCF21 | 37 | Female | Caucasian | CRS | 2 | Yes | 18 | 80 | No | Yes | No | No | Yes | GERD, Allergies | N/A |
| nonCF22 | 52 | Female | Caucasian | CRS | 1 | No | 5 | 20 | No | No | No | No | No | N/A | N/A |
| nonCF23 | 58 | Male | Caucasian | CRS | 0 | No | 11 | 25 | No | No | Yes | No | No | GERD, Allergies | N/A |

**Supplemental Table 1.** Clinical characteristics of CF (top) and non-CF (bottom) patients from whom nasal tissue was used, including Lund-Mackay (7, 8) and SNOT-22 (9) scores, both reflecting the severity of sinonasal disease symptoms. Abbreviations: Abx, history of antibiotics; AFS, allergic fungal sinusitis; ARS, allergic rhinosinusitis; CRS, chronic rhinosinusitis; DM, diabetes mellitus; FESS, functional endoscopic sinus surgery; GERD, gastroesophageal reflux disease; HTN, hypertension; N/A, not available; SNOT-22, 22 question sinonasal outcomes test.
